# A Newly Identified Role of the Tectorial Membrane in Aminoglycoside Ototoxicity

**DOI:** 10.64898/2026.05.04.722696

**Authors:** George W. S. Burwood, Pierre Hakizimana, Teresa Wilson, Rubing Xing, Wajiha Zaidi, Alfred L. Nuttall, Anders Fridberger

**Affiliations:** Oregon Hearing Research Center, Department of Otolaryngology – Head & Neck Surgery, Oregon Health & Science University, 3181 SW Sam Jackson Park Rd, Portland, OR, 97239, USA; Department of Biomedical and Clinical Sciences, Linköping University, Linköping, Sweden

## Abstract

Aminoglycoside (AG) antibiotic safety is limited by ototoxicity, the mitigation of which is vital considering bacterial resistance mediated erosion of our antibiotic arsenal. Previously, we observed tectorial membrane (TM) sequestration of Ca^2+^. We hypothesized that the TM sequesters other cations, including the AG gentamicin. We proposed to test the effect of TM genetic ablation on ototoxicity and TM-AG sequestration.

After intraperitoneal AG-furosemide, TM-lacking *Tecta*^ΔENT/ΔENT^ mice showed limited outer hair cell loss, unlike wildtype littermates. Spectroscopy measurements of gentamicin-Texas red (GTTR) were made in isolated wildtype and *Tecta*^Y1870C^ TMs and guinea pig cochleae following direct or intraperitoneal GTTR administration. TM-GTTR sequestration was observed in all cases, while negatively correlated with *Tecta*^Y1870C^ zygosity.

In summary, we discovered a novel TM component in the AG ototoxicity pathway. Intact TM structure is necessary for sequestration, and the TM modulates AG ototoxicity. TM-GTTR sequestration following systemic injection indicates that this phenomenon occurs during AG therapy.

**Single sentence summary:** Ototoxic aminoglycosides collect inside the acellular tectorial membrane of the inner ear, likely due to electrostatic interactions, and the structural status of that membrane modulates the toxic effect of those aminoglycosides on sensory hair cells.

## Introduction

Aminoglycoside antibiotics (AGs) are bactericidal agents which are highly effective in their treatment of Gram-negative bacterial infection. Gentamicin is still commonly used, for example in cases of premature infants who, if experiencing sepsis, may receive the drug for up to two weeks. AGs are experiencing a resurgence in use, partially in response to increasing bacterial resistance (1). However, a significant proportion of people treated with AGs experience ototoxic hearing loss. The prevalence of hearing loss following short course AG treatment (<16 days) is 16.6% of treated patients, translating to between 12.5 and 27.9 million people per year (2). Some 41% of patients receiving longer term AG treatment for tuberculosis develop hearing loss (3). Wider estimates of pediatric AG induced hearing loss indicate a maximum incidence of 61% (4). It is therefore evident that we maximize our understanding of AG hearing loss mechanisms and pathways.

AGs cross the blood labyrinth barrier into cochlear endolymph (5). The endolymph is a specialized, cytoplasm-like fluid that bathes the apical membranes of the hair cells. It is contiguous with the stria vascularis of the lateral wall, the structure that maintains the electrogenic endocochlear potential that underpins cochlear amplification (6, 7). The large potential difference across the sensory epithelium of the organ of Corti, from more than +110 mV in (murine) endolymph to -70 mV in the outer hair cells (OHCs) (e.g. (8)), is a major driving force behind cellular aggregation of AGs (at least acutely, see (9)). AGs predominantly enter the hair cells via the non-selective cation channel forming the pore of the mechanoelectrical transduction (MET) complex (10). Once taken up by OHCs, AGs are thought to induce production of reactive oxygen species, eventually triggering caspase activity and apoptosis (for review, see (11)). AGs can be found in organ of Corti sensory cells for up to 6 months after administration (12).

However, the hair cells of the organ of Corti are not directly exposed to endolymph. The tectorial membrane (TM), a collagenous extracellular matrix, is attached to the OHC stereocilia and sits closely to the inner hair cell (IHC) stereocilia, encapsulating the subtectorial space. The TM plays a crucial role in hearing, because it conveys the acoustic stimulus to the OHCs and provides a surface off which the reticular lamina may react – resulting in the shearing motion integral to MET channel activation and cochlear amplification (13, 14). Mutations to this structure can have a profound mechanical impact on hearing (15-21).

The role of the TM likely extends past its mechanical structure. Recently, it was demonstrated that calcium is sequestered by the TM to a concentration higher than the surrounding endolymph (22). Furthermore, sound exposure reduced this concentration, indicating that the sequestration is dynamic and contributes to normal cochlear function. The TM has negative fixed charge under physiological conditions (23) owing to a Donnan equilibrium (24). Since AGs are strongly polar and positively charged (25), we hypothesize that they are sequestered in the TM to a higher level than in surrounding endolymph, and that this reservoir of AGs influences hair cell ototoxicity. Increased concentrations of AGs in a structure in contact with the hair cells may be a neglected component of the ototoxicity process and thus could have implications for the interpretation of pathology, the design of dosing regimens and otoprotective protocols.

To address our hypothesis, we exposed mice lacking the protein α-tectorin, who thus lack a tectorial membrane, to acute ototoxic insult. We also employed fluorescence spectroscopy techniques in a variety of tissue preparations, using gentamicin Texas Red (GTTR) as our probe, to quantitatively explore the diffusion and concentration of AGs inside and outside of the TM. Isolated guinea pig temporal bones permitted examination of cochlear GTTR distributions in situ. Isolated murine TMs permitted examination of AG distribution in a controlled fluid environment, and genetic manipulation of the TM to identify protein/structural contributions to sequestration. Intraperitoneal injection of GTTR in mice allowed assessment of a clinically relevant question - do AGs sequester in the TM after systemic administration?

We found that mice lacking a TM lost susceptibility to AG induced OHC loss, and that guinea pig and mouse TM sequesters GTTR in situ, in vitro and in vivo. Truncation of α-tectorin in mice diminished sequestration capacity. We conclude that the TM is potentially a neglected component of AG ototoxicity.

## Results

### Mice lacking a TM are largely protected from ototoxic hair cell loss

We hypothesized that mice lacking a TM would be less sensitive to AG-induced ototoxicity. We IP injected mice with and without TMs with gentamicin and furosemide, in a protocol adapted from (26). We chose a dose of gentamicin and furosemide (GF) which would quickly produce a mild hair cell loss to avoid the potential ceiling effect of extensive hair cell loss. *Tecta*^+/+^ and *Tecta*^ΔENT/ΔENT^ littermates were injected with 200 mg/kg of gentamicin followed by 200 mg/kg of furosemide 30 minutes later. They were monitored, given soft food and heat support for 4 hours in the laboratory, as the furosemide induced reduced activity, likely due to blood pressure drop. We monitored animal weight and administered supportive saline, and provided soft food in the vivarium. Animals were euthanized and cochleae harvested, fixed and decalcified at the 72 hour timepoint, as strial defects become apparent after this timepoint in similar protocols (27) and we aimed to avoid significant strial contributions to the ototoxicity. Cochleae were dissected and wholemounts were prepared for OHC counts.

Fig. 1 shows the results of the functional ototoxicity assay. Fig. 1A shows example cochlear wholemounts from *Tecta*^+/+^ and *Tecta*^ΔENT/ENT^ mice following GF treatment. Notably, the OHCs in the hook regions of the GF treated *Tecta*^ΔENT/ENT^ animals were intact. *Tecta*^ΔENT/ΔENT^ mice treated with furosemide only generally had intact hair cell populations.

**Figure 1:**
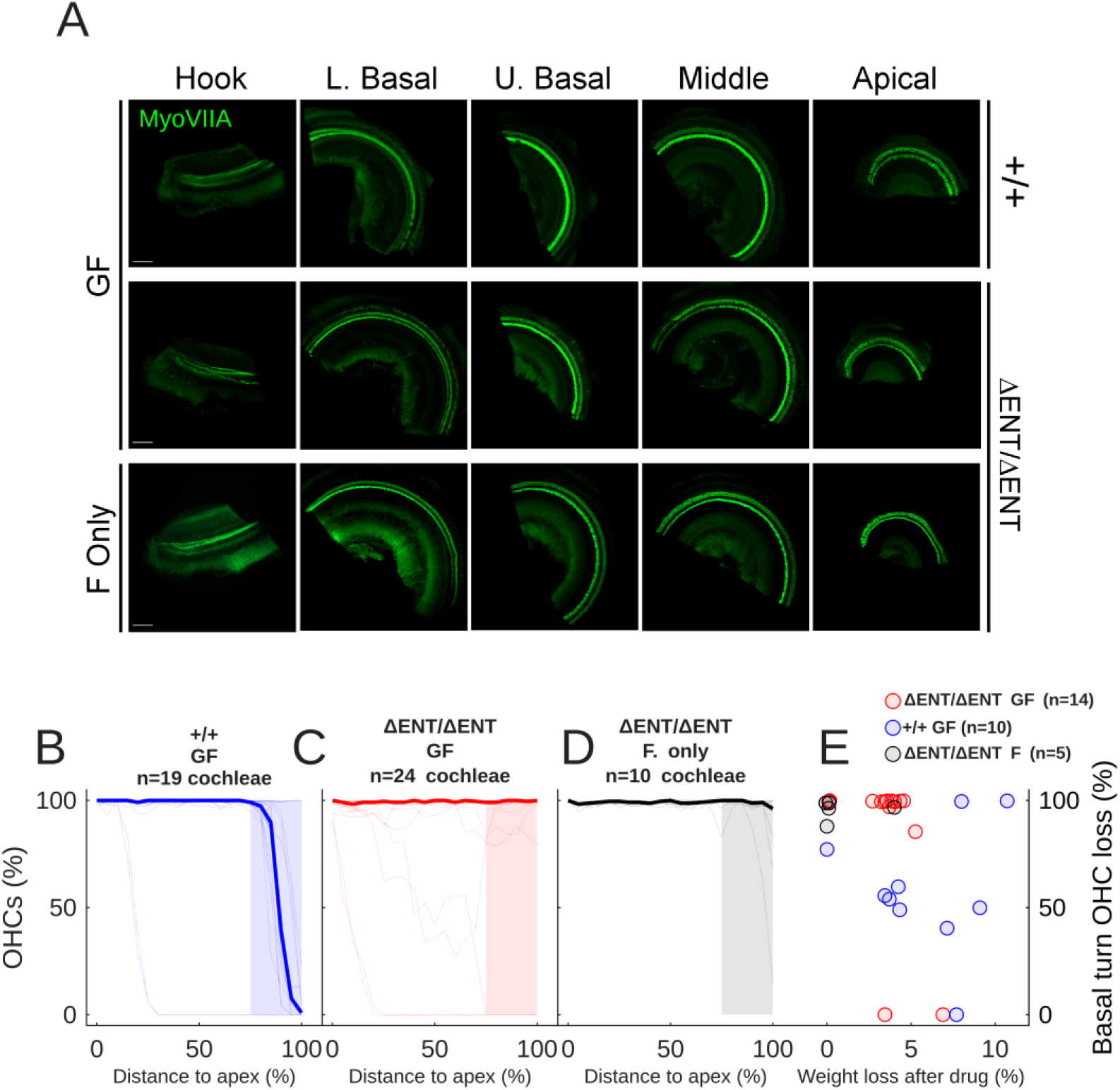
A: Example organ of Corti wholemount preparations from two *Tecta*^ΔENT/ΔENT^ mice, one treated with GF and one only treated with F, and one GF treated Tecta^+/+^ mouse, immunolabelled with Myosin VIIA. Scale bar 150 μm. B: Percentage of surviving OHCs as a function of cochlear distance to the apex in 19 cochleae (blue lines, median shown by thick blue line) from 10 Tecta+/+ mice 72h after they were injected IP with 200 mg/kg gentamicin and 200 mg/kg furosemide (GF). C: As in B, but for 24 cochleae from 14 *Tecta*^ΔENT/ΔENT^ mice (red lines, median shown by thick red line). D: As in C but for a group of 10 cochleae from 5 *Tecta*^ΔENT/ΔENT^ mice treated with furosemide only (black lines, median shown by thick black line). E: OHC survival in the shaded areas of B-D as a function of peak percentage weight loss following GF injection, for 14 *Tecta*^ΔENT/ΔENT^ mice (red circles) and 10 Tecta^+/+^ littermates (blue circles), and 5 *Tecta*^ΔENT/ΔENT^ mice treated with F only.

The OHC loss was quantified in 10 *Tecta*^+/+^ mice (Fig. 1B). 3/10 mice showed no OHC loss – this is an expected outcome as the GF protocol has been previously shown to be only partially effective. 6 of 10 mice showed only hook-lower basal turn OHC loss, while 1 mouse showed extensive OHC loss. The basal turn first loss profile is typical as basal turn OHCs are more vulnerable to multiple types of insult. The median OHC loss (blue thick line) indicates that the typical OHC loss region was restricted to 80-100% of longitudinal distance to the apex.

The above-mentioned basal turn OHC loss was not the typical outcome for *Tecta*^ΔENT/ΔENT^ mice. Fig. 1C shows the OHC loss profiles of the *Tecta*^ΔENT/ΔENT^ mice. Only 2 of 14 mice displayed extensive OHC loss, and only 1 of those 2 mice showed the expected basal-apical pattern of loss – and it is notable that the mice with the most loss in both groups were from the same litter. The remaining *Tecta*^ΔENT/ΔENT^ mice did not display OHC loss or showed mild, sporadic loss. The median OHC loss (red thick line) indicates that the typical outcome was a lack of OHC loss.

A single dose of furosemide alone is not expected to induce OHC loss in normal animals (28, 29). To control for the possibility that hair cell loss may occur due to the *Tecta*^ΔENT/ΔENT^ mutation, we injected 5 *Tecta*^ΔENT/ΔENT^ mice with furosemide only (Fig. 1D). The results show that only 1 cochlea displayed severe hair cell loss in the lower basal turn, out of the 10 analyzed (there was no OHC loss in the other cochlea from this animal), demonstrating that most of the effect seen in wildtype animals was due to the GF insult.

Hair cell loss severity was significantly dependent upon genotype (repeated measures ANOVA adjusted for multiple comparisons: Genotype:Hair cell loss between 75 and 100% of the distance from the apex (shaded areas, Fig. 1B-D): p=4.17×10^-5^). Post-hoc testing (Tukey) showed significant differences between GF treated genotypes at 95 and 100% of the distance from the apex (p=0.0044 and p=0.00029 respectively), but not for the lower frequency regions, and not between GF and F only treatment groups.

To confirm that there was not a relationship between OHC loss severity and weight loss (e.g. due to dehydration or variability in dose absorption), we plotted the correlation between percentage peak weight loss following drug and the amount of hair cell loss within the shaded areas of Fig. 1 B-D (Fig. 1E). It was typical for mice to lose some body weight in the 72 hours following drug injection. However, weight loss did not contribute to OHC loss severity (repeated measures ANOVA adjusted for multiple comparisons: p=0.404).

In summary, genetic ablation of the TM led to reduced OHC toxicity and occasional disruption of the typical tonotopic axis of OHC loss after ototoxic insult. These data suggest that the TM plays a functional role in outer hair cell ototoxicity in response to AGs.

### The in vitro murine TM sequesters GTTR and *Tecta* is necessary for sequestration efficacy

Having observed that the TM modulates AG ototoxicity, we investigated the interaction between the TM and AGs in vitro. We placed freshly isolated mouse TMs in artificial endolymph (AE) containing 26 uM calcium chloride. We performed fluorescence correlation spectroscopy (FCS) measurements of the movement of GTTR inside the TMs of both CBA/CaJ mice and each genotype of the *Tecta*^Y1870C^ line. FCS is a statistical technique whereby the stoachastic light emission of low concentrations of a fluorescent dye is recorded over time as they diffuse through a confocal detection volume. The resulting time series of spikes can be autocorrelated and fitted to a diffusion model (see methods), to derive the amount, speed and behavior of that dye, depending on the experimental parameters.

We hypothesized that TM GTTR concentrations will be higher, and speed of diffusion will be lower, in intact TMs, and mutation of the TM will decrease concentration and increase diffusion speed. In the temporal bone preparation, dye would continuously diffuse away from the measurement site, and only very small amounts of dye were added locally. By measuring from isolated TMs under in vitro conditions, we could tightly control and homogenize the GTTR concentration in the AE,thus using it as a ratiometric reference.

Fig. 2A-C shows transmitted light images of the lateral edge of the murine TM. Parallel fibers are visible in the wildtype TM (Fig.1A). The pattern of the parallel fibers appears partially disrupted and more wrinkled in the heterozygous animal (Fig.1B). Finally, the parallel fibers are disrupted and the structure is diffuse in the homozygous mutant (Fig.1C).

**Figure 2:**
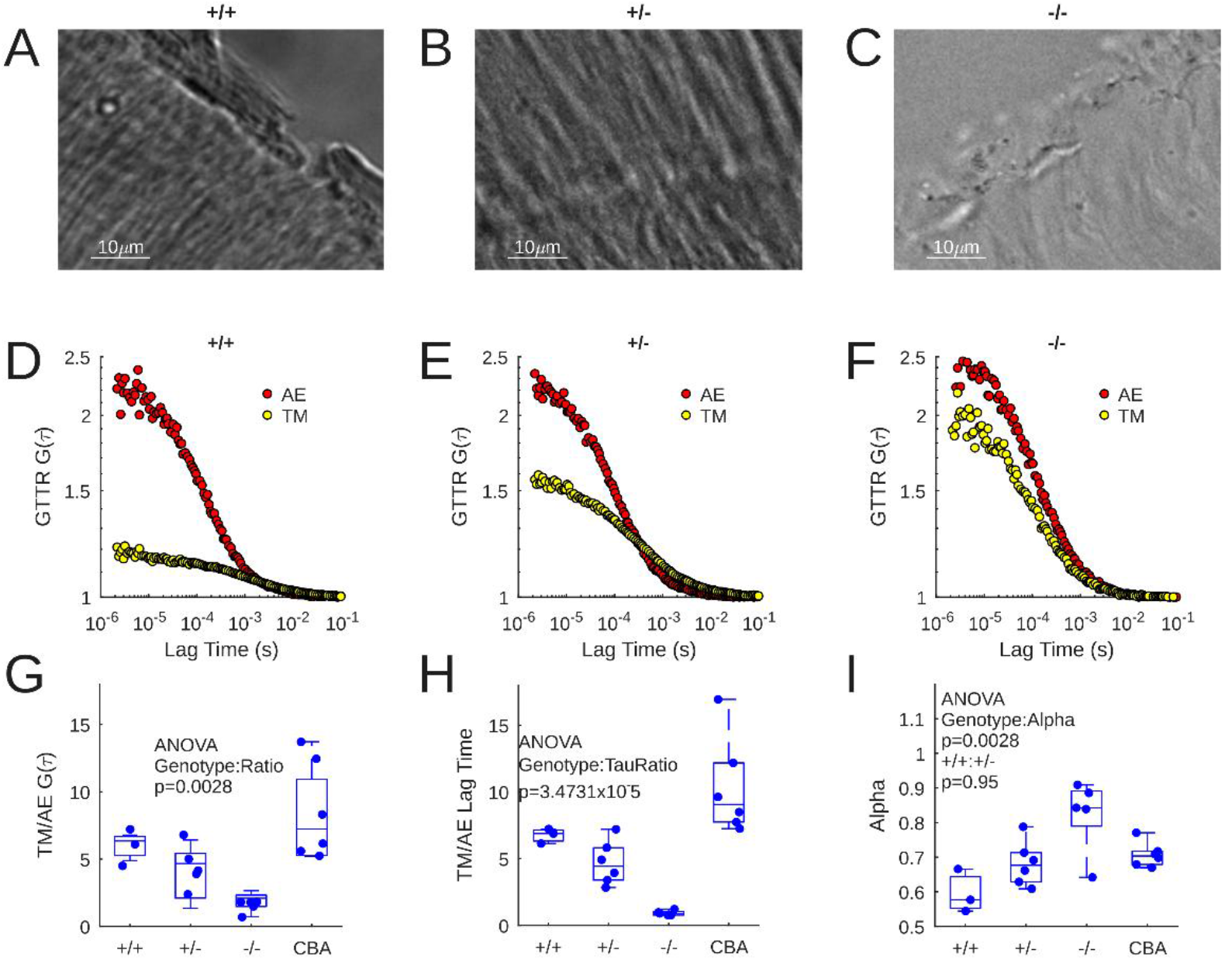
A-C: 63x transmitted light micrograph of a wildtype TM (A), a *Tecta*^Y1870C/+^ TM (B), and a *Tecta*^Y1870C/Y1870C^ TM (C). D-F: Example autocorrelation functions from AE (red) and TM (yellow) for wildtype (D), *Tecta*^Y1870C/+^ (E), and *Tecta*^Y1870C/Y1870C^ (F) experiments. G: Ratio of TM and AE G(τ) values for 3 *Tecta*^+/+^, 6 *Tecta*^Y1870C/+^, 6 *Tecta*^Y1870C/Y1870C^, and 6 CBA/CaJ experiments. H: TM/AE lag time ratio for the experiments shown in G. I: Alpha value for the experiments shown in G and H.

The correlation function G(τ) corresponds to the number of GTTR molecules within the confocal detection volume, where the closer the initial G(τ) value is to 1, the higher the number of molecules. Example autocorrelation functions from AE (red dots) and the TM (yellow dots) are shown for CBA/CaJ, *Tecta*^Y1870C/+^ and *Tecta*^Y1870C/Y1870C^ derived samples (Fig. 2D-F). The largest difference between autocorrelation curves can be seen in the CBA/CaJ records (Fig. 2D), implying that there are more GTTR molecules inside the TM than in the surrounding AE. The *Tecta*^Y1870C/+^ curve differences are smaller (Fig. 2E), and smaller still for *Tecta*^Y1870C/Y1870C^ (Fig. 2F).

These data are statistically summarized in Fig. 2G. GTTR sequestration was expressed as a ratio of G(τ) in the TM versus that of surrounding endolymph. FCS performed in 6 isolated CBA/CaJ TMs mice demonstrated TM/AE ratios that were significantly larger than that measured in the *Tecta*^Y1870C/Y1870C^ (p=0.0020). The TM/AE ratio was not statistically different between *Tecta*^+/+^ and CBA/CaJ (p=0.5323). However, the TM/AE ratio of the *Tecta*^+/+^ mouse was not significantly different from the other groups. One way ANOVA indicated that genotype had a significant effect on TM/AE ratio (p=0.0028). This means that intact TMs buffer GTTR at concentrations on average five-fold higher than surrounding fluid, and sometimes over ten-fold.

The lag time of the correlation function, which corresponds to molecular diffusion speed, was also examined between genotypes (Fig. 2H). The TM/AE ratio of lag time was significantly different for CBA/CaJ and both *Tecta*^Y1870C/+ &^ *Tecta*^Y1870C/Y1870C^ (p=0.0024 and > 0.0001 respectively). The lag time ratio was significantly different for *Tecta*^+/+^ and *Tecta*^Y1870C/Y1870C^ (p=0.0131). Overall, lag time ratio was significantly affected by genotype (ANOVA, p=3.4731×10^-5^). This means that more intact TMs constrain GTTR diffusion.

The anomalous diffusion model used to fit the FCS data derived from the TMs includes a term, alpha, which is an estimate for the barriers to diffusion. In free fluid, Alpha is 1 implying Brownian motion, and decreases as diffusion becomes more anomalous. We compared the Alpha values across genotype (Fig. 2I) – there was a statistically significant impact of genotype upon Alpha (ANOVA p=0.0028). The mean Alpha of the *Tecta*^Y1870C/Y1870C^ was significantly higher than those of *Tecta*^+/+^ and *Tecta*^Y1870C/+^ (p=0.0022 and 0.0208 respectively). Alpha derived from CBA/CaJ measurements was not statistically different from other groups. This means that intact TMs appear to induce anomalous, constrained diffusion that is slower than that of Brownian motion.

### GTTR sequestration in the TM is not spatially homogeneous

We correctly hypothesized that mutation of the TM would reduce its sequestration capacity and speed up GTTR diffusion. However, FCS is sensitive to noise from aggregates, debris or background fluorescence, and point-measurement position within the TM may contribute to variability within the dataset.

To further assess this issue, we also used a spectroscopy technique, Number & Brightness analysis (N&B), that was more robust to noise and that gave more information on the spatial behavior of GTTR sequestration inside the TM.

N&B scans were made from isolated TMs from the 4 mouse strains described above, in identical conditions. Fig. 3 shows the N&B findings. Fig. 3A-C show the average molecular number in each pixel of a 128×128 scan of the TM repeated 300 times, for *Tecta*^Y1870C/+^ (Fig. 3A), *Tecta*^Y1870C/Y1870C^ (Fig 3B) and CBA/CaJ (Fig. 3C). The colormap is scaled to Fig. 3C. Fig. 3 A’-C’ show the molecular brightness for the scans in A-C. The TM and the surrounding AE are labelled. The *Tecta*^Y1870C/+^ TM in Fig. 3A shows some sequestration, but a relatively disrupted fibrous structure with several gaps – which appear closer in color to the AE. These gaps could influence FCS measurements. For the *Tecta*^Y1870C/Y1870C^ TM in Fig. 3B, the Number of the TM structure and that of the AE are comparable. For the normal TM in Fig. 3C, there is substantial difference between the Number in the TM and AE, indicating sequestration capacity. The brightest part of the image is on the interface between the AE and the TM – possibly at the marginal band.

**Figure 3:**
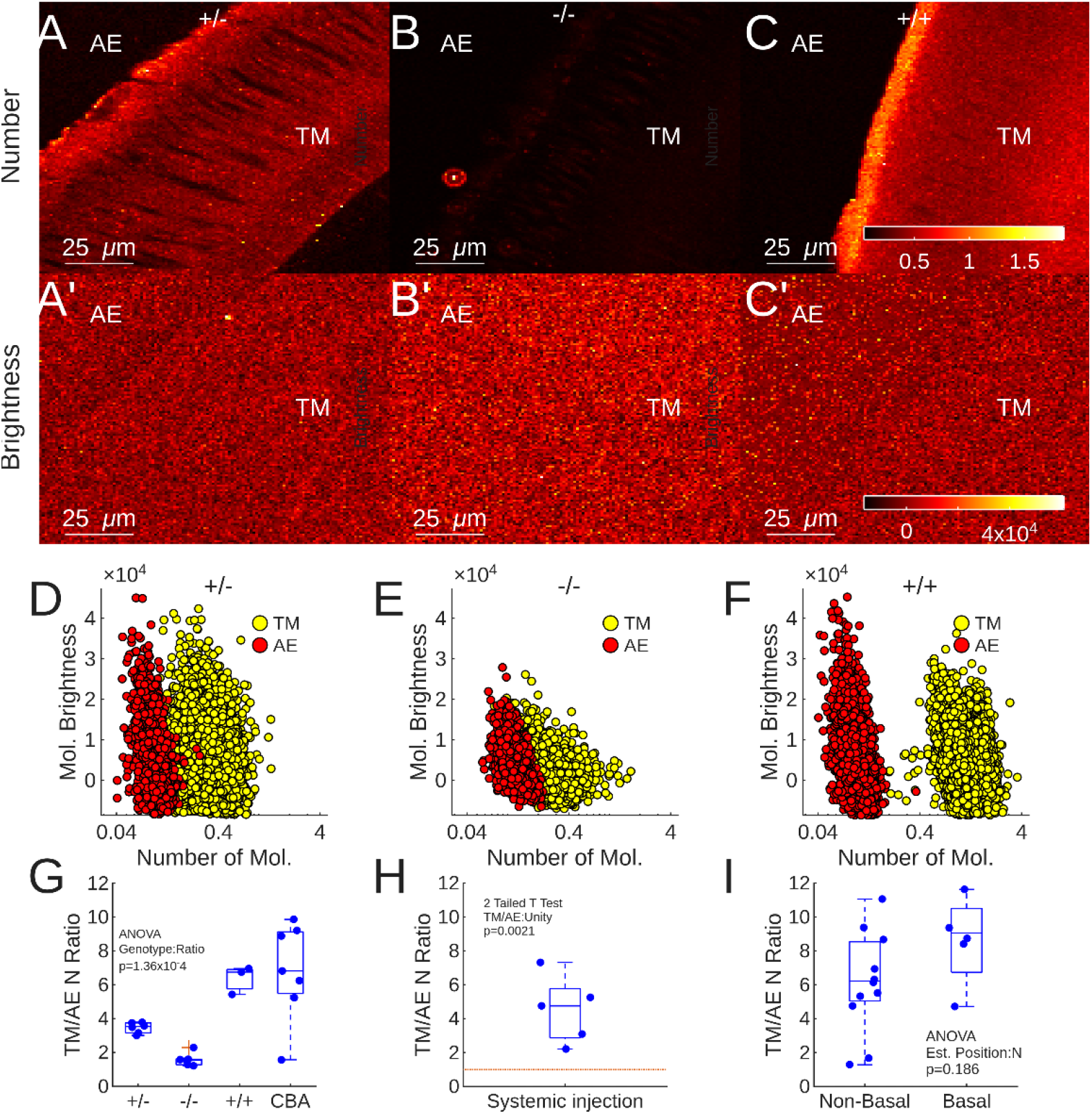
A-C: Mean number of molecules present in *Tecta*^Y1870C/+^ (A), *Tecta*^Y1870C/Y1870C^ (B), and CBA/CaJ (C) TMs. The AE and TM portions of the image are labelled. A’-C’: Mean molecular brightness for the TMs in A-C. D-F: N&B plots for the images in A-C and A’-C’. G: Quantified TM/AE number of molecules in several preparations (CBA/CaJ N=7, *Tecta*^+/+^ N=3, *Tecta*^Y1870C/+^ N=6, *Tecta*^Y1870C/Y1870C^ N=5). H: TM/AE number of molecules ratio in 5 *Tecta*^+/+^ mice euthanized 2 hours after systemic injection with 0.5 mg/kg GTTR. I: TM/AE number of molecules ratio in non-basal and basal TM samples from both *Tecta*^+/+^ and CBA/CaJ mice.

The homogeneous nature of the images in A’-C’ indicate that the location of the GTTR, or the TM genotype, did not influence its molecular brightness.

Pixel number & brightness scatter plots are shown in Fig. 2D-F for each of the example scans in Fig. 3A-C’. The number of GTTR molecules appears marginally higher in the TM (yellow dots) relative to AE (red dots), for both *Tecta*^Y1870C/+ &^ *Tecta*^Y1870C/Y1870C^ while the molecular brightness distributions are similar for both areas (Fig. 3D-E respectively). For the CBA/CaJ TM, there is separation between the TM and AE clusters indicating a higher number of GTTR molecules inside the TM.

To quantify the data shown in Fig. 3D-F and control for variations in dye concentration between experiments, the ratio between the molecular number in TM and AE is expressed as a function of genotype (Fig. 3G). ANOVA demonstrated that number of molecule ratio is highly dependent upon genotype (p=1.36×10^-4^. CBA/CaJ N=7, *Tecta*^+/+^ N=3, *Tecta*^Y1870C/+^ N=6, *Tecta*^Y1870C/Y1870C^ N=5). Both CBA/CaJ and *Tecta*^+/+^ TMs demonstrate a sequestration capacity, that is not statistically different (p=0.9802) while both having significantly higher sequestration capacity than that recorded from *Tecta*^Y1870C/Y1870C^ TMs (p=0.0002 & 0.0044 respectively). The statistical significance between both CBA/CaJ and *Tecta*^+/+^ and *Tecta*^Y1870C/+^ is less compelling (p=0.0110 & p=0.1082 respectively). These data confirm the FCS findings that the normal TM sequesters gentamicin and that sequestration is disrupted by mutation of *Tecta*.

### GTTR accumulates in the murine TM after IP injection

In addition to innoculating the TM with GTTR locally, we wished to demonstrate that GTTR, and thus GT, sequesters inside the TM following systemic administration of the drug. We injected wildtype mice with 0.5 mg/kg of GTTR in sterile saline. After 2 hours, the animals were euthanized, and the TMs were harvested for correlation spectroscopy imaging. No GTTR was added to the bath. Fig. 3H shows that the ratio of the TM/AE number of molecules is significantly greater than 1 (T test, p=0.0021, N=5). This experiment confirms that systemically injected GTTR accumulates inside the TM, suggesting that this may occur in the clinic. It should be noted that the ability to calculate a TM/AE ratio implies that GTTR could exit the TM into the AE – thus suggesting it could act as a source of AGs and not just a sink.

### Longitudinal differences in TM AG sequestration capacity are not apparent in this dataset

The TM possesses multiple indices of longitudinal gradients, such as cross-sectional area, stiffness and mass. These gradients arise from the structural proteins of the TM, and we hypothesized that they may also result in gradients of molecular sequestration capacity which could result in different concentrations of AGs across the tonotopic gradient.

To test this hypothesis, we separated the harvested murine TM pieces into groups, where those pieces thought to be from the apex or middle turn (non-basal) were compared with those originating from the basal turn (basal), from CBA/CaJ and *Tecta*^+/+^ mice. Since the dissection of the fresh TM often resulted in its fracturing and only parts of it being imaged, it was not possible to obtain precise longitudinal location data. The TM/AE number ratios are shown in Fig. 3I. While there appears to be a higher ratio for pieces identified as basal, the difference was not statistically different (ANOVA, p=0.186).

### The guinea pig TM sequesters GTTR in situ

It is possible that species differences lead to different AG-TM interaction characteristics. To test the hypothesis that TM AG sequestration occurs independent of species, and to assess the behavior of GTTR in the intact organ of Corti, we performed FCS on the organ of Corti of our guinea pig temporal bone preparation. Note that these experiments were performed using a different optical setup to those in mouse, due to limitations imposed by using the intact preparation. Thus, perform FCS with a lens of lower numerical aperture (1.1 versus 1.41) and magnification (32x versus 63x), leading to less spatially restricted measurements.

Following decapitation, guinea pig temporal bones were quickly dissected, placed in a custom holder, the cochlea was exposed and the apical turn opened in artificial perilymph. GTTR was administered through the intact Reissner’s membrane via micropipette. Fig. 1 shows the absolute number of GTTR molecules detected outside and inside the TM immediately after injection.

Fig. 4A shows a laser scanning micrograph measuring the fluorescence emission of tissue in the absence of GTTR. Small concentrations of endogenous fluorescence are visible, outlining the surface of the reticular lamina and supporting cells. Neither TM nor hair cell morphology are visible. A nanomolar concentration of GTTR was introduced to the scala media via picospritzer driven micropipette. The distribution of GTTR fluorescence in the same location as Fig. 4A is then shown in Fig. 4B. The TM can now be distinguished from the endolymph. Subdomains of the TM, closer to the IHC and OHC region can be distinguished. The stereocilia of the IHCs and OHCs are now strongly fluorescent, as well as the cell somae – indicating ongoing uptake of the GTTR molecule.

**Figure 4:**
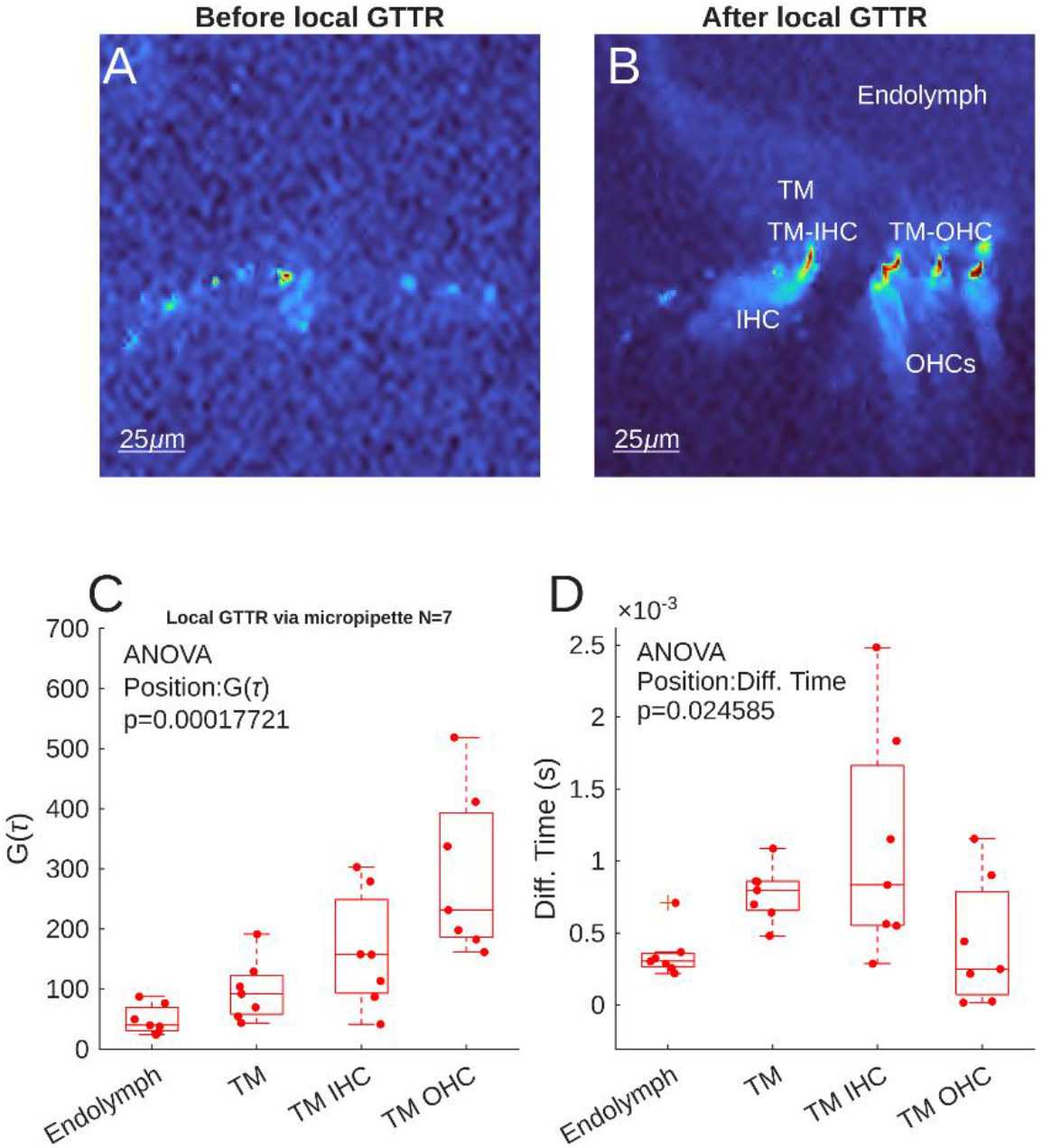
A:Background fluorescence of the guinea pig apical organ of Corti prior to local GTTR injection. B: the same apical organ of Corti following GTTR injection. The Endolymph can be distinguished from the TM, and TM-IHC and TM-OHC domains are apparent, as well as the IHCs and OHCs (labelled). C: Quantification of the effective number, G(τ), of GTTR molecules present in each location in 7 preparations. D: The diffusion time of those GTTR molecules in the same measurements as C.

The locations identified in Fig. 4B were then used as targets for FCS. Quantification of FCS data from 7 preparations are shown in Fig. 4C. Following injection, the effective number of GTTR molecules in endolymph was 49.23 ±24.12. Inside the central body of the TM, there was a statistically non-significant increase in GTTR molecule number (97.75 ± 50.78).

Close to the IHCs inside the TM, the mean effective number of GTTR molecules was 162 ± 96.71. The highest mean GTTR molecule number was found inside the TM, close to the OHCs, at 291.55 ± 143.88. There was a larger variation in the number in the hair cell adjacent TM microdomains. Overall, there was a highly statistically significant relationship between measurement position and GTTR molecule number (ANOVA, p=0.00018).

Fig. 4D shows the diffusion times associated with the population data in Fig. 4C. There was a significant relationship between location and diffusion time (ANOVA, p=0.025). Diffusion time in the main body of the TM and TM-IHC location was higher (0.8±0.19 ms and 1.1±0.79 ms respectively) than that of endolymph (0.4±0.16 ms, p=0.36 and p=0.031 respectively) – implying an anomalous, confined diffusion behavior, rather than Brownian motion, was statistically present close to the IHCs. However, the mean diffusion time was lower in the TM-OHC location (0.4±0.43 ms), and not statistically different from endolymph (p=0.99). This is possibly due to the sink effect of residual OHC MET current, or local structural differences e.g. Kimura’s membrane.

Overall, these data show that locally applied GTTR can sequester within the guinea pig TM to levels higher than endolymph, and there is possibly a microdomain effect due to proximity to the hair cells.

## Discussion

We have demonstrated that the ototoxicity of gentamicin is modulated by the presence or absence of the TM in vivo (Fig. 1), the TM of guinea pigs and mice can accumulate AGs (Fig. 2-4), that this accumulation occurs after systemic injection (Fig. 3), is disrupted following mutation of α-tectorin in mice (Fig. 2, 3,), and that. The core finding of TM AG sequestration is significant as the TMs role in ototoxicity has been neglected.

We speculate that the reduction in the number of calcium binding sites in the Y1870C variant of *Tecta* is partially responsible for the loss of TM AG sequestration in these mutants. As the striated sheet matrix is highly disrupted in homozygous Y1870C mutants (as indicated by a diffusion time, Alpha and lag time value more similar to those expected from fluid, see Fig. 2), meaning that GTTR molecules “inside the TM” may be only weakly affected by their environment. The Y1870C mutation reduces fixed charge (from -6.4 mmol/l to -2.1 mmol/l) (30), and the proportionality of the fixed charge reduction approximates the reduction in AG sequestration seen here in *Tecta*^Y1870C/+^ mutants relative to wildtypes (Fig. 2G, 3G). Thus, our data support the notion that α-tectorin is a contributor to TM fixed charge and therefore is likely a major driver of AG sequestration. It is, however, unclear whether the calcium binding domains of the peptide structure itself, or the charge of its glycoconjugates, is the main driver of this mechanism.

Our attempts to assess longitudinal sequestration differences were not successful, mainly because we did not precisely measure longitudinal position (Fig. 3I). As stated above, fresh, wildtype TMs were fragile and often broke into pieces. Attempts were made to image each end of long pieces, but the overall difference did not turn out to be statistically significant. *Tecta*^Y1870C/+^ TMs were more likely to be dissected intact – probably due to their weaker association with the organ of Corti. *Tecta*^Y1870C/Y1870C^ TMs were difficult to handle as they often adhered to the Reissner’s membrane and were extremely fragile. The longitudinal gradient of TM AG sequestration is thus an open question which requires further study, but it is tempting to speculate that at least part of the positive apex to base AG vulnerability gradient (e.g. Fig. 1B) is related to TM sequestration capacity.

The experiments in Fig. 1 suggest that the intact TM contributes to ototoxicity, and the spectroscopy experiments suggest that the increased local concentration of AGs in the TM also plays a role. This finding chimes with the observation that sound exposure reduces TM calcium (22) and increases OHC calcium (31). This suggests that the TM acts as a local source for positively charged molecules, and that interactions between the MET channel’s electric field and the fixed charge of the TM could be influenced by pharmacology, and homeostasis disrupting events such as loud sound exposure (32). Through this interaction, the presence of AG in the TM may modulate the MET channel activity, and thus the rate of AG uptake by hair cells.

The spatial distribution of AGs in the scala media has been shown to be different than expected, due to its accumulation in the TM. This is significant for understanding of ototoxicity mechanisms because we do not currently know the half-life of TM-AG interaction. While serum levels of AGs may reduce in hours - 3 hours for GTTR versus 56 minutes for unconjugated gentamicin (33) - the TM may sequester AGs for longer than this elimination period. It has been shown that concentration of gentamicin in perilymph increases between 3 and 5 hours after systemic administration (34). Gentamicin has been detected via immunohistochemical assay in multiple inner ear tissues, including OHCs at 6 months following a single administration of drug in guinea pigs, although images were only provided for these cells at the 3 month mark (12). The TM could act as a long-term source of AGs, which, during a repeated dosing regime, is continuously refreshed. It is also known that AGs can buffer long-term in the stria vascularis. AG potentiation of noise induced hearing loss has been shown to occur within 20 days of drug administration (35), and latent AG presence in the TM and/or the stria could contribute to this effect (5).

Concerning the impact of AGs upon the lateral wall, recent research shows that gentamicin may induce TNF-α mediated apoptosis of lateral wall pericytes in a murine chronic AG treatment model (36). Interruption of this pathway led to hair cell loss mitigation. This observation points to the possibility that loss of pericytes may lead to gradually higher cochlear penetration of AGs over the course of chronic treatment, due to gradually worsening dysfunction of the blood labyrinth barrier. Evidence for this occurrence could be found in the TM – we speculate that chronic AG treatment leads to gradually increasing TM AG sequestration. Nonetheless, our use of a rapid AG-diuretic treatment paradigm highlights the role of the TM in high frequency outer hair cell loss while minimizing strial contributions. This, and the incomplete mitigation of hair cell loss by TNF-α pathway inhibition, suggests that protection of strial integrity may not be sufficient to prevent AG induced hearing loss without addressing TM sequestration.

Another factor to consider is whether the TM regulates the open probability of the OHC MET channels. Mice lacking a TM may have been shown to have a lower MET channel open probability and less MET channel activity overall due to the lack of TM-hair cell shear displacement (17). The lack of displacement is important as even relatively mild sound exposure induces movement of calcium from the TM to the OHCs (22). Additionally, endocytosis is a far slower process than MET channel activity (10). When TM sequestration is absent, AGs may be cleared more rapidly from the scala media, and so OHCs may less efficiently take them up due to both shortened exposure time and reduced MET open probability. Data to the contrary regarding the TM’s regulation of MET operating point exist (37). Under these conditions, one might expect sequestration to be the dominant component influencing ototoxicity outcomes in the gentamicin/furosemide experiments we performed. Further experiments involving MET channel mutant strains crossed with TM null mutants could be performed to answer this question using AGs.

In summary, we have identified a novel component of the ototoxicity mechanism in the mammalian inner ear. This mechanism, based upon TM sequestration of positively charged molecules (22), may contribute to the time course of AG ototoxicity by acting as a drug sink directly adjacent to the highly vulnerable OHCs. This sequestration may potentially contribute to the differing ototoxicity effects of other small molecule drugs such as cisplatin (38-40), which is predicted to become cationic only after cellular entry (41), thus making it unlikely that it would sequester in the TM. The role of the TM in ototoxicity should be further studied, with a view to mitigating clinically induced ototoxicity and improving the efficacy and safety profiles of ototoxic drugs, including AGs.

## Acknowledgements

The authors would like to thank the Oregon Health & Science University Advanced Light Microscopy Core (RRID: SCR_009961) for their assistance with FCS and N&B equipment. We thank Dr. Mary Ann Cheatham for the provision of *Tecta*^Y1870C^ mice, Dr. Guy Richardson for helpful advice, and MRC Harwell, UK, for the provision of cryopreserved *Tecta*^ΔENT^ spermatozoa. Funding support was provided by the National Institute on Deafness and Communication Disorders at the National Institute of Health (NIH R01-DC000141 (Nuttall, Fridberger), Swedish Research Council (2022-00548, Fridberger), Swedish Brain Foundation (FO2023-0171, Fridberger).

## Author contributions (CRediT)

GB: Conceptualization, Funding acquisition, Methodology, Investigation, Data Curation, Formal Analysis, Visualization, Supervision, Project Administration, Writing (original draft), Writing (review & editing).

PH: Investigation, Methodology, Writing (review & Editing).

TW: Methodology, Investigation, Formal Analysis, Visualization, Project administration, Writing (original draft), Writing (review & editing).

RX: Investigation, Project administration, Writing (review & editing). WZ: Investigation.

AN: Conceptualization, Funding acquisition, Supervision, Writing (review & editing).

AF: Conceptualization, Funding acquisition, Methodology, Investigation, Data Curation, Formal analysis, Project administration, Resources, Software, Supervision, Validation, Writing (review & editing).

## Methods

### Animals

All procedures were conducted in accordance with approved OHSU IACUC protocols (IP00001278) and with the consent of the local ethics committee. Mice of both sexes and various ages were used for FCS and N&B experiments, but all mice were under 6 months old. CBA/CaJ mice (Jackson Labs, 000654) were initially used in this study while breeding of *Tecta*^Y1870C^ mice was taking place. *Tecta*^Y1870C^ mice were kindly provided by Dr. Mary Ann Cheatham, and were backcrossed onto the CBA/CaJ line for 8 generations. *Tecta*^ΔENT^ mutant mouse sperm were procured from MRC Harwell, UK and rederived onto a CBA background and backcrossed for several generations before starting ototoxicity experiments. For ototoxicity experiments, mice were between 11 and 12 weeks of age. Procedures on young guinea pigs were carried out with approval of the Regional Ethics Committee in Linköping, Sweden (permit 19058/2021).

### TM dissection

Mice were euthanized with an overdose of ketamine and xylazine and then decapitated. The cochleae were extracted and placed in AE (in mM: 148 KCL, 1.3 NaCl, 5 HEPES, 3 Glucose, with 26 μM CaCl_2_, pH 7.3, 300 mOsm/kg) in a dish under a dissecting microscope. Excess tissue such as the stapedial artery, fascia and muscle was removed with tweezers, and the boney wall of the face of the cochlea was removed using bent tweezers. Once as much of the bone as possible was removed, the modiolus was sectioned at the base using tweezers. The cochlear spiral was then gently prised out of the remaining otic bone. This step could result in damage to the organ of Corti, but the TM often remained attached to the inner sulcus.

The cochlear spiral was then held at the modiolus end with tweezers, and a sterilized eyelash was used to separate the TM from the spiral. Most of the time, up to half a turn was salvaged, but sometimes it was possible to extract most of the apical-middle turn TM. Often, however, the hook region was lost. The tip of a tungsten microelectrode was also used to remove TMs, but this led to more breakage.

The free pieces of TM were then located and moved via a BSA coated 20 ul pipette to a fresh AE bath. The BSA helped prevent the TM from sticking to the tip, and only the very last of the pipette’s draw was used to move the TM so it did not travel too far into the tip.

Finally, the TM pieces were deposited into a glass bottomed dish that had been coated with Cell-Tak (Corning). The TM was gently manipulated and pressed down onto the Cell-Tak using an eyelash, so that it did not move when the dish was agitated. Dishes were then placed on ice. AE volume was typically 20ul per TM piece and was recorded.

### GTTR preparation

GTTR was purchased from AAT Bioquest (24300) and diluted to a 100uM stock solution in AE. For FCS and N&B imaging, working solutions of 1uM were prepared in AE and thoroughly vortexed to reduce dye aggregation, before being stored on ice and in darkness.

### Ex-vivo guinea pig temporal bone preparation

7 Young guinea pigs weighing less than 400 g were deeply anesthetized with intraperitoneal sodium pentobarbital and decapitated. After decapitation, the temporal bones were isolated and placed in a plastic holder. The bulla was opened, but the middle ear was kept intact, allowing sound stimulation through a loudspeaker positioned in the ear canal. The preparations were kept viable by immersion in tissue culture medium and by perfusing oxygenated tissue culture medium through scala vestibuli after making small openings at the cochlear base and apex. The apical cochlear opening was also used to position a glass microelectrode in scala media, which was used to deliver GTTR solution using a picospritzer. The apical opening was also used for confocal imaging and FCS with a Nikon 40x water immersion lens with numerical aperture 0.8.

### FCS & N&B imaging

Spectroscopy was performed on a Zeiss LSM980 inverted microscope in photon counting mode, managed by the OHSU Advanced Light Microscopy Core, using a 63x oil immersion lens N.A. 1.4. After placing the TM sample dish on the stage, the pieces of TM were located using the 10x lens. Background FCS recordings were made in AE and inside the TM. GTTR was then added to the dish for a final concentration of 20-100nM, and the final volume was usually made up to be 500ul. For systemic injection experiments, no background recordings were possible. FCS point scans were made in 10 10s repetitions, or 50 2s repetitions if encountering substantial noise. Laser power was typically 0.2%, wavelength was 561 nm, the pinhole was 63 um. The same parameters were used within experiments to allow for direct comparison of fluid and TM recording ratios. The detector bandwidth was opened widely above the stimulation wavelength.

N&B imaging was normally conducted following FCS measurements. The lens was focused on the marginal edge of the TM in order to include TM and AE regions of interest. A 128×128 square image was recorded in raster mode with 300 repetitions, with either 32 or 16 us pixel dwell time.

## FCS analysis

FCS data were analyzed using custom MATLAB software. Individual averages were recorded, and so noisy correlation curves could be excluded from the analysis. Curves were deemed to be noisy if they contained excessive undulation, evidence of bleaching, or poor correlation at long lag times. Background recordings from AE and TM were subtracted from FCS recordings as appropriate.

Autocorrelation functions (G(τ)) were fitted with a diffusion with triplet model in the case of AE:

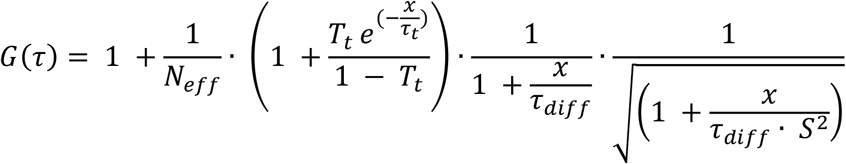

where χ is the lag time, N_eff_ is the effective number of molecules, T_t_ is the triplet function, τ_t_ is the triplet lifetime,

τ_diff_ is the diffusion time, and S is the axial/lateral ratio of the illumination volume. and an anomalous diffusion with triplet model in the case of TM:

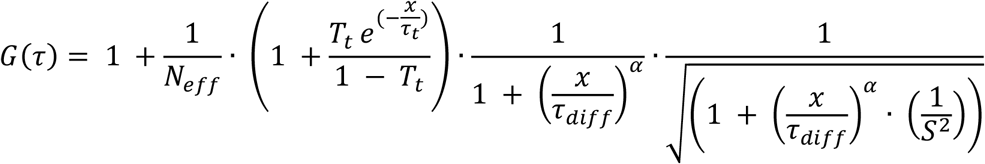

Where α is the anomalous diffusion exponent representing the constrained environment of the TM.

Corrected effective number of molecules, diffusion lag time, and alpha values (in the case of anomalous diffusion) were recorded, ensuring that model parameters were not fixed at bounds and the R^2^ was at least 0.990.

The Corrected effective number of molecules and lag time were then treated as ratios between the TM and AE to account for differences in fluorescence/imaging conditions. Alpha values were reported raw.

### N&B analysis

N&B data were also analyzed using custom MATLAB software. Images were opened using bioformats (Mathworks) and regions of interest were drawn on the TM and inside the AE. A boxcar average was applied to the full time course of the fluorescence data to mitigate the effect of bleaching.

### Systemic GTTR injections

Mice were IP injected with 0.5 mg/kg GTTR in sterile saline. After the injection, the mice were placed in their home cage on a warming pad for two hours, and provided with soft food. After 2 hours, the mice were sacrificed and cochleae harvested as described above. TMs were kept in dark and on ice after dissection, and underwent N&B imaging as described above without further addition of GTTR.

### Ototoxicity Protocol

*Tecta*_ΔENT/ΔENT_ and wildtype littermates were injected IP with 200mg/kg Gentamicin (G1264, Sigma Aldrich) in sterile saline followed by 200mg/kg furosemide (71288-203-04, Meitheal Pharmaceuticals) 30 minutes later. Following injections, the animals were monitored until they started to recover from the acute effects of furosemide which could include lethargy and reduced activity – up to a maximum of 4 hours. Animals were provided with soft food. Following monitoring, animals were returned to the vivarium and weighed daily. If they had lost weight, they were administered with 1 ml warmed saline IP. At 72 hours post drug treatment, animals were euthanized and the cochleae were harvested, fixed and decalcified. Confocal imaging of cochlear wholemounts was conducted, using Myosin7A (1:300, 25-6790, Proteus Biosciences) to identify hair cells, using a protocol adapted from (42, 43).

### Statistical analysis

Data are reported means ± standard deviations. ANOVA, Repeated Measures ANOVA, Linear Mixed Models and Student’s T tests were used. The formula for the linear mixed model was *Hair Cell Count* ∼ *Genotype* × *Weight loss* + (1|*Distance from Apex*). Significance was assumed at a p value of less than 0.05 or adjusted as appropriate for multiple comparisons.

## Notes

### Competing Interest Statement

The authors have declared no competing interest.

